# Guiding Irregular Nuclear Morphology on Nanopillar Array for Malignancy Differentiation in Tumor cells

**DOI:** 10.1101/2022.01.30.478168

**Authors:** Yongpeng Zeng, Yinyin Zhuang, Aninda Mitra, Peng Chen, Isabella Saggio, G. V. Shivashankar, Weibo Gao, Wenting Zhao

## Abstract

For more than a century, abnormal nuclei in tumor cells, presenting subnuclear invaginations and folds on the nuclear envelope, have been known to be associated with high malignancy and poor prognosis. However, current nuclear morphology analysis focuses on the features of the entire nucleus, overlooking the malignancy-related subnuclear features in nanometer scale. The main technical challenge is to probe such tiny and randomly distributed features inside cells. We here employ nanopillar arrays to guide subnuclear features into ordered patterns enabling their quantification as a strong indicator of cell malignancy. Both breast and liver cancer cells were validated, as well as the quantification of nuclear abnormality heterogeneity. The alterations of subnuclear patterns were also explored as effective readouts for drug treatment. We envision this nanopillar-enabled quantification of subnuclear abnormal features in tumor cells opens a new angle in characterizing malignant cells and studying the unique nuclear biology in cancer.

**Teaser:** A nanopillar-based assay quantifying the abnormal nuclear morphology in tumor cells at single-cell level.

## Introduction

Nuclear polymorphism is one characteristic feature widely reported across a variety of cancer types (*1*). However, clinical grading of tumor cell’s nuclei for pathological diagnosis still largely relies on subjective visual inspection of nuclear morphology through optical microscopic imaging (*2*), which counts on the experience of individual pathologists and inevitably suffers from poor reproducibility and reliability (*3*). To address this bottleneck, a variety of technologies, including clinical sample labeling (*4*), computer-aided medical image processing with machine learning or artificial intelligence (*5*, *6*), and microfabrication and micropatterning for the whole nucleus guidance (*7*, *8*), have been explored for objective and quantitative characterization of the nuclear architecture. Nevertheless, the quantifiable parameters are mainly focused on nuclear size and circularity, uniformity of nuclear chromatin, karyoplasmic ratio, and nucleoli, etc. (*5*, *9*); while the obvious subnuclear abnormalities including folds, invaginations, and inclusions on nuclear envelopes are only evaluated qualitatively due to their random appearance in individual nuclei and huge variations between cells. More critically, a significant technical challenge for their quantification lies in their near-diffraction-limited size (i.e. in tens to hundreds of nanometers scale), which is hard to characterize under optical microscopy.

To probe subcellular properties near or below diffraction-limited size scales, vertically aligned nanopillar arrays have recently been introduced as a powerful tool. They were shown to effectively interrogate plasma membrane properties at the scale of tens to hundreds of nanometers, such as altering the membrane permeability for intracellular delivery of biomolecules (*10*–*13*), performing subcellular electroporation and recording intracellular electrophysiology signals (*14*, *15*), manipulating nanoscale membrane topography to recruit endocytic proteins (*16*, *17*) and trigger actin polymerization (*18*), probing cellular mechanics (*19*, *20*), and guiding cell adhesion, migration and differentiation (*21*, *22*). More interestingly, nanopillars have been reported to reach the nucleus in the intracellular space for probing nuclear deformability and their cytoskeleton regulators in live cells (*23*), altering the distribution of different lamin proteins along them (*24*), and rewiring the mechanotransduction from the plasma membrane to the nucleus (*24*). However, whether nanopillar-induced nuclear deformation correlated with the subnuclear irregularities in cancer cells and how such correlation can be used to assess cancer cell properties have not been explored before.

In this study, we demonstrate a quantitative characterization of subnuclear irregularity in cancer cells using vertically aligned nanopillar arrays. When plated on nanopillars, the subnuclear features of cancer cells are effectively guided into ordered deformation patterns with readable anisotropicity. Interestingly, the increase of anisotropicity shows an obvious correlation with higher malignancy and faster cell migration. Taking advantage of the single cell resolution with multiple sampling points per nucleus, we evaluate both the heterogeneity in a cancer cell population and the differential response of anti-metastatic drugs between high and low malignant cells.

## Results

### Nanopillar guides nuclear shape irregularities in cancer cells

Differential response of subnuclear morphology to nanopillars among tumor cells with different malignancies was first evaluated (Fig. 1A). Arrays of vertically aligned nanopillars (as shown in the scanning electron micrograph (SEM) (Fig. 1B) were fabricated on transparent quartz coverslip using electron-beam lithography (EBL) and reactive ion etching (RIE). Taking breast cancer cells as a model, we examined two well-known cell lines that exhibit distinct metastatic potential, low malignant MCF-7 and high malignant MDA-MB-231 cells, on both flat and nanopillar arrays. After an overnight culture to allow sufficient generation and stabilization of deformations on nanopillars, the nuclear morphology was visualized via immunostaining of nuclear lamina protein lamin A, a key modulator of nuclear shape in pathogenesis (*25*). For low malignant MCF-7 cells, they display a smooth nuclear outline on flat substrates without noticeable subnuclear features (representative image shown in Fig. 1C top left panel and more examples shown in fig. S1A left column). In contrast, those cultured on nanopillar arrays generate an ordered array of lamin A rings colocalizing with nanopillar position underneath (Fig. 1C bottom left panel and fig. S1A right column), consistent with the previous report (*23*). In comparison, the high malignant MDA-MB-231 cells, on the flat surface, show prominent but randomly distributed subnuclear foldings and wrinkles across the nucleus, indicating altered nuclear architecture yet challenging to quantify with irregular shapes (Fig. 1C top right panel and fig. S1B left column). But when cultured on nanopillar arrays, MDA-MB-231 cells exhibit significantly decreased randomness of subnuclear irregular features. Instead, distinct alignment of subnuclear features into line patterns along adjacent nanopillars are clearly observed (Fig. 1C bottom right panel and fig. S1B right column), suggesting a remodeling of the lamin network guided by the local perturbations from nanopillars.

**Figure 1.**
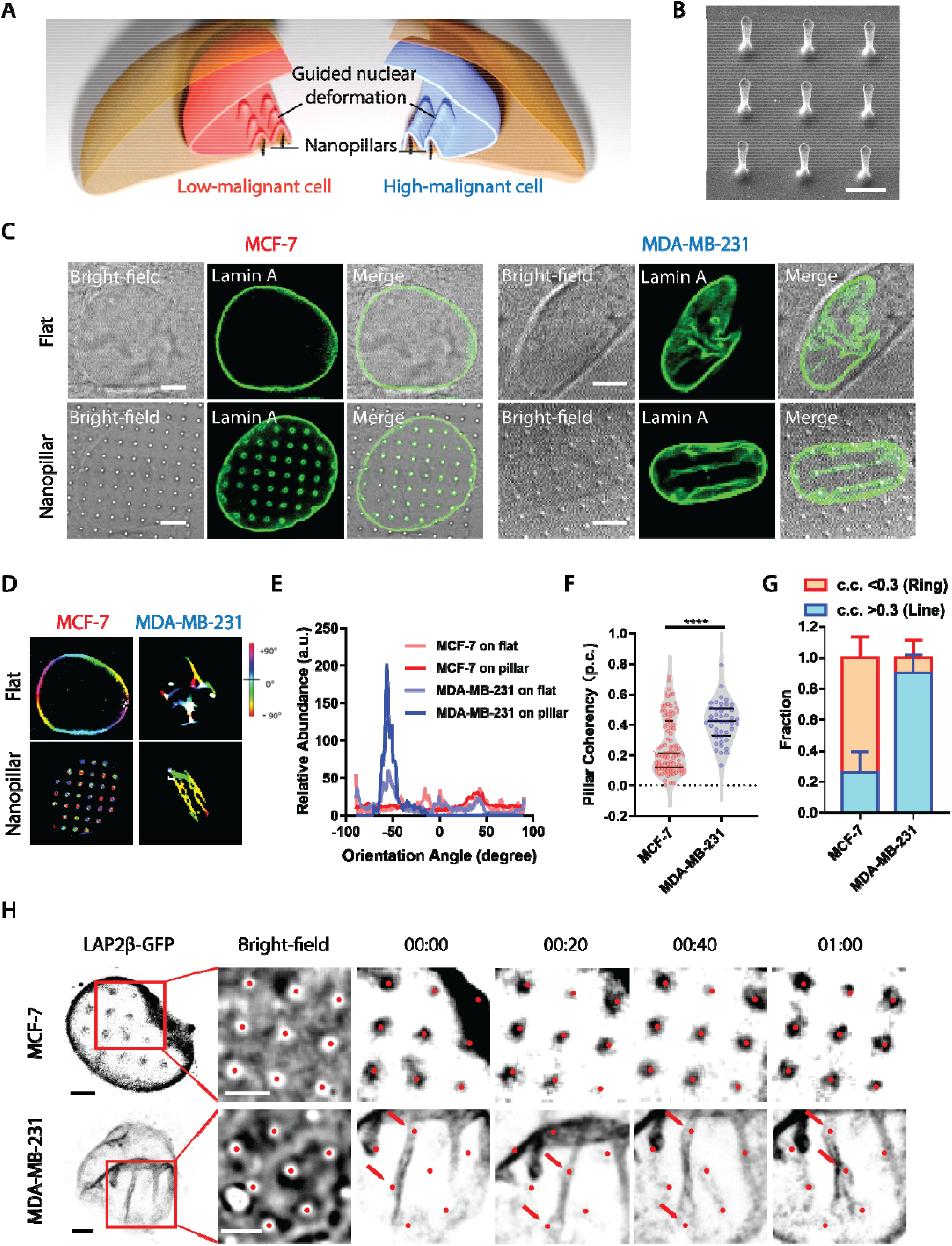
Nanopillar-guided subnuclear deformation patterns correlate with cancer malignancy. (**A**) Schematics of different patterns generated by nanopillar-guided nuclear shape deformation in cancer cells with varying malignancies. (**B**) SEM of nanopillar arrays. Scale bar, 2 μm. (**C**) Nuclear morphology of MCF-7 cells and MDA-MB-231 cells on a flat surface. Scale bars, 5 μm. (**D**) Orientation of nuclear shape irregularities and nanopillar-guided nuclear features in MDA-MB-231 cells and MCF-7 cells. (**E**) Comparison of orientation distribution of nuclear shape irregularities and nanopillar-guided nuclear features in MDA-MB-231 cells and MCF-7 cells. (**F**) Anisotropy measurement of the nanopillar-guided nuclear deformation in MCF-7 cells (N = 94 pillars) and MDA-MB-231 cells (N = 44 pillars). (**G**) Fraction of ring deformation and line deformation in MCF-7 cells (N = 39 cells) and MDA-MB-231 cells (N = 24 cells). Ring deformation is defined by c.c. < 0.3 whereas ring deformation is defined by c.c. > 0.3. (**H**) Dynamics of nanopillar-guided nuclear features in MCF-7 and MDA-MB-231 cells for one hour. Red dots indicate nanopillar locations. Red arrows in the bottom row refer to the nanopillars that guide the nuclear grooves. Scale bars, 3 μm. Statistical significance of the two groups was compared using an unpaired t test with Welch’s correction, p-value: ****<0.0001.

Based on the guided subnuclear features on nanopillars, a quantitative analysis of lamin A is performed to differentiate MCF-7 and MDA-MB-231 cells. One pronounced difference is the isotropicity of the lamin A network generated patterns on nanopillars: the rings formed in MCF-7 nuclei give isotropic distribution around nanopillars, while the lines aligned across pillar arrays in MDA-MB-231 cells generate anisotropic intensity profile around each nanopillar with dominant distribution in certain angles. To quantitatively distinguish the isotropic and anisotropic patterns of subnuclear deformation on nanopillars, we removed the nuclear boundary and analyzed specifically the orientation distribution of the lamin A’s subnuclear patterns using the OrientationJ (*26*) plug-in in ImageJ (*27*), where for each pixel, the angle that aligns dominantly with surrounding signals is calculated and displayed in color hues. The intact images with nuclear boundaries of example cells in Fig. 1C for each condition are shown in fig. S2, where the nuclear boundary of MCF-7 on a flat surface remains uncut as no subnuclear feature is detectable. As shown in Fig. 1D, the isotropic ring deformation of MCF-7 nuclei on nanopillars exhibits a broad and random angle distribution of subnuclear features and thereby generates mixed colors surrounding each nanopillar (Fig. 1D, bottom left). Similarly, the nuclear boundary of MCF-7 on the flat surface also displayed a combination of different angles in individual nuclei, and thus a variety of colors (Fig. 1D, top left). In contrast, the aligned subnuclear deformation in MDA-MB-231 on nanopillars gives rise to a preferred angle across the whole nucleus and thus displaying a prominent yellowish color (Fig. 1D, bottom right). However, such a pillar-guided nuclear pattern in high malignant MDA-MB-231 cells could not be formulated in flat surfaces, which only showed a random distribution of subnuclear groves and invaginations (Fig. 1D, top right). Here, by collecting the angle distribution inside the individual nucleus, we found that the nanopillar array produces a detectable predominant angle for the anisotropic subnuclear deformation in high malignant MDA-MB-231 cells. In contrast, no dominant subnuclear orientation is observed in the low malignant MCF-7 nucleus (Fig. 1E).

For quantitative evaluation of deformation orientations, the anisotropy of the subnuclear features on each nanopillar is further converted into orientation coherency values (i.e., pillar coherency, or p.c. in short), which is ranging from 0 to 1 with 0 representing a completely isotropic pattern (e.g. perfect circle) and 1 refers to an extremely anisotropic pattern (e.g. straight line). Low malignant MCF-7 cells have much lower coherency value (p.c.=0.27 ± 0.18, n=94 pillars) than high malignant MDA-MB-231 cells (p.c.=0.42 ± 0.13, n=44 pillars) (Fig. 1F). By averaging the p.c. values of the same cell, we can further obtain an averaged cell coherency value (in short as c.c.) as a single cell readout for cell population analysis. Based on the statistics on individual pillars, we set a coherency value of 0.3 as the threshold to distinguish isotropic and anisotropic subnuclear features in this study. Not surprisingly, MCF-7 contains a higher fraction of ring-deformation cells (c.c. < 0.3, fraction=0.74 ± 0.13, n=39 cells) while MDA-MB-231 sample mainly contains line-deformation cells (c.c. > 0.3, fraction=0.90 ± 0.11, n=24 cells), as shown in Fig. 1G. It is interesting to note that both low-malignant MCF-7 cells and high-malignant MDA-MB-231 cells contain a mixed population of low c.c. and high c.c. cells, indicating a possible heterogeneity of canner nuclear properties even within the same cell type. In addition, we validate such nanopillar-guided nuclear deformation in two typical liver cancer cells with divergent invasiveness, highly invasive SK-HEP-1 and non-invasive PLC-PRF-5 (*28*). By wound healing assay, we found that invasive SK-HEP-1 cells migrated faster than the non-invasive PLC-PRF-5 cells (fig. S3A). Interestingly, when both cell lines were cultured on nanopillar arrays, invasive SK-HEP-1 cells displayed anisotropic line patterns formed in the nuclei, whereas non-invasive PLC-PRF-5 cells exhibited isotropic ring patterns on nanopillar arrays (fig. S3B). The distribution of their p.c. values shows significant difference between these two cell lines (fig. S3C), and the c.c. values effectively differentiate the two cell lines apart (fig. S3D). Taken together, these results confirmed that nanopillar arrays can effectively guide subnuclear morphological irregularities in tumor cells and can generate quantifiable subnuclear readouts for cell malignancy evaluation with single-cell resolution.

In addition, the dynamics of such subnuclear features on nanopillar are further examined in live cells via transient expression of nuclear envelope protein, LAP2□ fused with green fluorescent protein (LAP2□-GFP). Strikingly, subnuclear rings in MCF-7 cells are relatively stable on each nanopillar location over 1 hour regardless of the overall movement of the whole nucleus (Fig. 1H upper row), while aligned line patterns of LAP2□-GFP in MDA-MB-231 cells are able to switch among nearby nanopillar sites following the migration of nucleus (Fig. 1H lower row). It confirms that nanopillars provide persistent guidance on subnuclear deformations and serve as stable sampling points for subnuclear irregularity measurement.

### Probe cancer heterogeneity via guided nuclear deformation

Heterogeneity among cancer cells presents one of the major hurdles in cancer therapy, as cancer metastasis or drug resistance are often observed only in a subset of cells with higher malignancies (*29*). Taking the advantages of the single-cell resolution of this nanopillar-based nucleus grading system, we were motivated to verify its accuracy in characterizing the heterogeneity of nuclear irregularity in a mixed cell population. First, we generate a series of cell mixtures containing cancer cells of different malignancies by mixing GFP-CAAX tagged low malignant MCF-7 cells with unlabeled high malignant MDA-MB-231 cells in predefined ratios (illustrated in Fig. 2A). A typical microscopy image of a cell mixture containing both cell types on nanopillar arrays is shown in Fig. 2B, where both line deformation in an unlabeled MDA-MB-231cell and ring deformation in GFP labeled MCF-7 cells were observed. When plotting the fraction of cells with high deformation coherency (>0.3) on nanopillars against the ratio of unlabeled MDA-MB-231 in the cell mixture, we observed a positive correlation between nanopillar-measured fraction and the portion of high malignant MDA-MB-231 cells in the mixture (Fig. 2C). It indicates that cancer cells in heterogeneous malignancy can be sorted and quantitatively characterized via the nanopillar-guided subnuclear deformation patterns.

**Figure 2.**
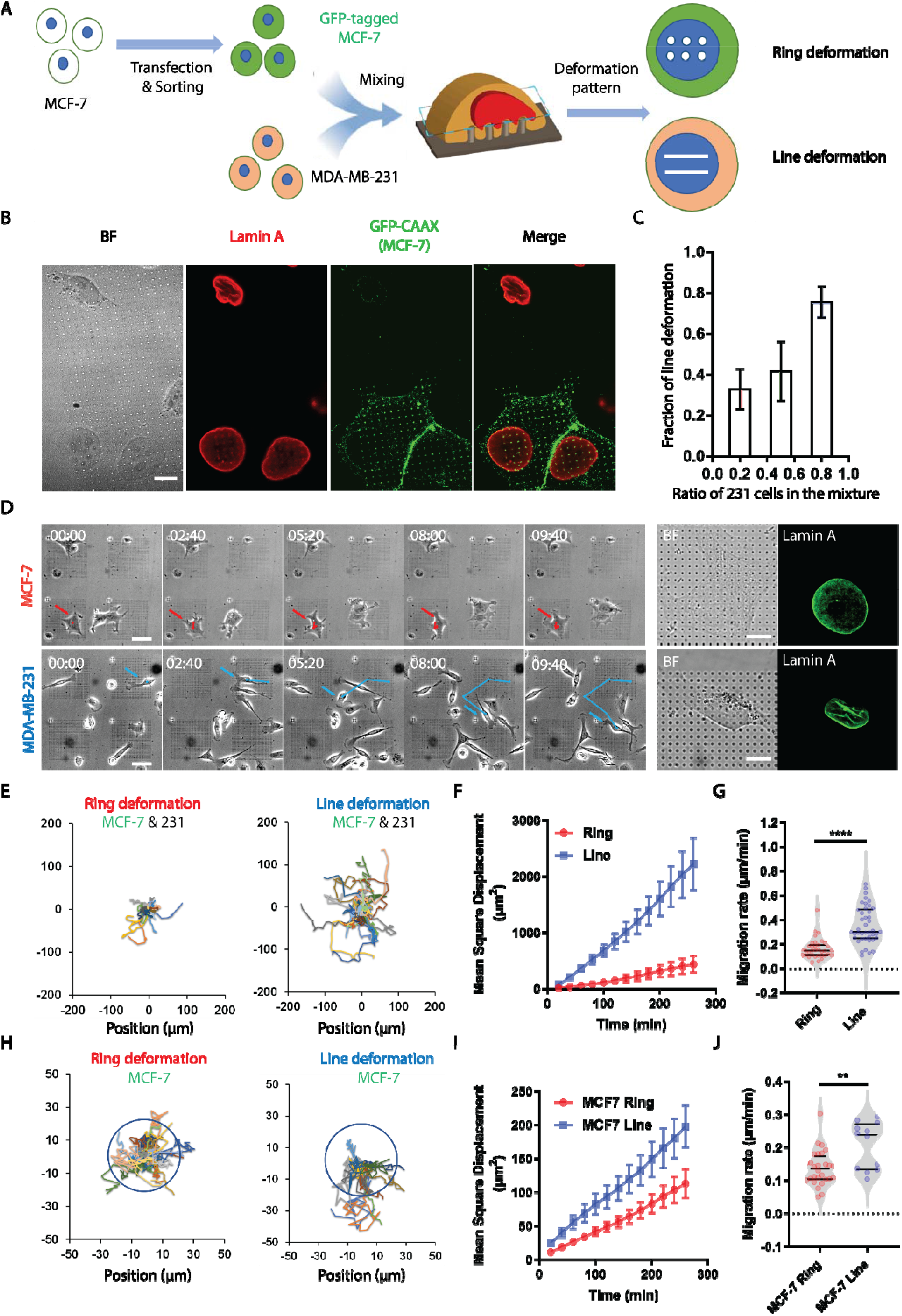
Probing cancer heterogeneity via nanopillar-induced subnuclear deformation. (**A**) Methodology for using nanopillar arrays to probe cancer heterogeneity. (**B**) MDA-MB-231 cell and GFP-tagged MCF-7 cells showing different nuclear deformation patterns on the same substrate. Scale bar 10 μm. (**C**) Correlation between fraction of cells showing line-like guided nuclear deformation and the ratio of MDA-MB-231 cells to MCF-7 cells. (**D**) Brightfield images of MCF-7 cells and MDA-MB-231 cells migrating on nanopillar arrays over time and the fluorescent images showing deformation patterns of cells at the last time point. Scale bars for left figures, 50 μm. Scale bars for right figures, 10 μm. (**E**) Migration trajectories of cells showing different nuclear deformation patterns on nanopillars (Ring: N= 35 cells; Line: N= 42 cells). (**F**) MSD measurement of cells showing different nuclear deformation patterns. (**G**) Comparison of migration rate of cells showing varying nuclear deformation patterns. (**H**) Migration trajectories of MCF-7 cells showing different nuclear deformation patterns on nanopillars (MCF7 ring: N= 28 cells; MCF7 line: N= 12 cells). Two blue circles with the same diameter are centered with the origin to show that MCF-7 cells with line deformation on nanopillars tend to migrate faster than those showing ring deformation. (**I**) MSD measurement of MCF-7 cells showing different nuclear deformation patterns. (**J**) Comparison of migration rate of MCF-7 cells showing varying nuclear deformation patterns. Statistical significance of migration rate measurement under different conditions was evaluated by an unpaired t-test with Welch’s correction. ****P < 0.0001; **P < 0.01

In addition, nanopillar-remodeled lamin A patterns are also strongly correlated with cell migration speed, another key indicator of malignant cells. Cell motility of individual cells obtained from live-cell imaging was correlated with their subnuclear deformation patterns on nanopillars for both MCF-7 and MDA-MB-231 cells (Fig. 2D). When pooling together both cells with same nanopillar-remodeled patterns (ring or line separately), we found that lamin A patterns are correlated with cell migration speed by having ring-formation in slow-migrating cells and line-forming within fast-migrating cells despite their cell types, which are clearly shown in cell migration trajectories (Fig. 2E), mean square displacement (MSD) (Fig. 2F), and calculated cell migration rate (Fig. 2G). More interestingly, even among all the low malignant MCF-7 cells, two subpopulations with distinct migrating speed can be differentiated based on nanopillar-guided lamin A patterns (Fig. 2, H-J, and fig. S4). MCF-7 cells with line deformation migrate faster than MCF-7 cells with ring patterns on nanopillars, as similarly measured by longer migration trajectories (Fig. 2H), larger mean square displacement (MSD) (Fig. 2I), and faster cell migration rate (Fig. 2J). It is plausible to speculate that the polarized contractility in fast migrating cells contribute to the high coherency of aligned lamin A patterns. Altogether, the nanostructured platform constitutes an effective technology for probing the heterogeneity in a cancer cell population with single cell resolution.

### Evaluate anti-metastatic drug effect via guided nuclear deformation

Given the ability to identify and characterize high malignant or fast migrating cancer cells in a mixed cell population, we next sought to use the established nanopillar sensing system to evaluate anti-metastatic drugs. More than 90 percent of cancer mortality is caused by cancer metastasis (*30*). Identifying and developing ant-cancer drugs, particularly those specifically targeting metastasis-prone cells, is an emerging route for new cancer therapy (*31*). However, the development of anti-metastasis drugs is a daunting yet challenging task as metastasis only develops from a subset of cells and is difficult to evaluate using conventional methods probing the whole cell population, such as western blot, transwell migration, and matrigel invasion assays (*32*). In comparison, nanopillar-guided subnuclear deformation effectively identifies high malignant or fast migrating cancer cells with single-cell resolution through lamin A line pattern. Therefore, we hypothesize that the conversion of line patterns of high malignant cells in response to drug treatment can be used to evaluate drug effectiveness against metastasis (as illustrated in Fig. 3A). As a proof of concept, we examined a reported anti-metastatic drug, curcumin (*33*, *34*), and compared the deformation changes of MCF-7 and MDA-MB-231 on nanopillars in response to it. As shown in Fig. 3B, the high-malignant MDA-MB-231 cells exhibited significantly less line deformation, while low-malignant MCF7 showed no significant pattern changes. Upon quantification, both decreased pillar coherency values (DMSO, 0.42 ± 0.13, n=34 pillars; curcumin, 0.24 ± 0.16, n=52 pillars) (Fig. 3C) and decreased fractions of line-deformation cells (DMSO, 0.93 ± 0.12, n=27 cells; curcumin, 0.26 ± 0.27, n=30 cells) (Fig. 3D) are observed for MDA-MB-231 after curcumin treatment. No significant change in the anisotropy of induced nuclear deformation is found in MCF-7 cells (Fig. 3, C and D). Consistently, MCF-7 cells do not show significant changes in migration speed (p value=0.7440) in response to the curcumin treatment, while MDA-MB-231 cells exhibit a lower migration rate (p value=0.0363) (Fig. 3, E and F) after the same treatment. Concentration dependency was further characterized, where both the pillar coherency value and fraction of line-deformation cells responded sensitively to as low as 1 μM (fig. S5). Besides curcumin, we also evaluated another reported anti-metastatic drug, haloperidol (*35*), and obtained similar responses as shown in fig. S6, which further validated that the anisotropy of nanopillar-guided subnuclear deformation can be an effective indicator for anti-metastatic drug evaluation.

**Figure 3.**
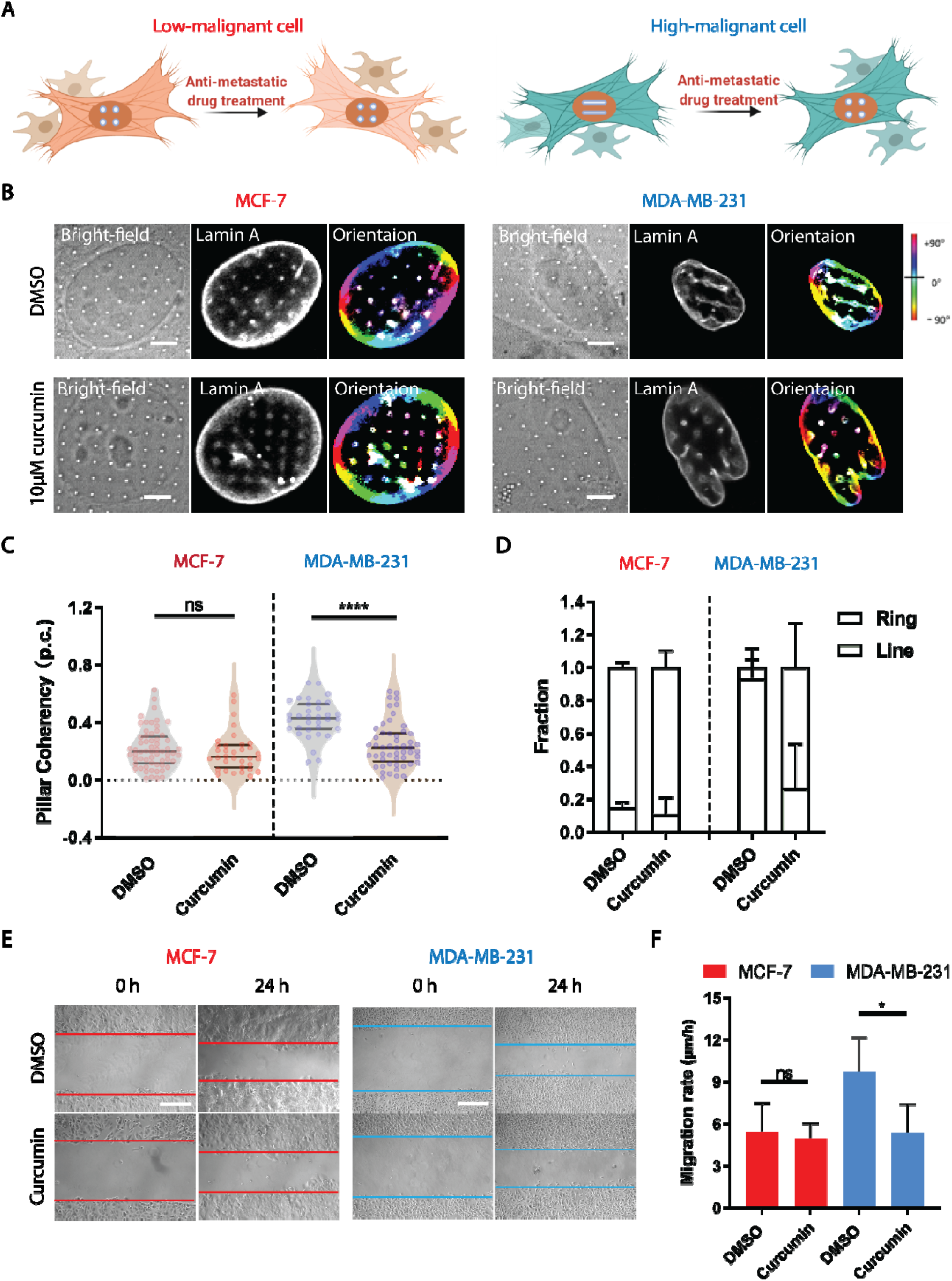
Evaluating anti-metastatic drug effects via nanopillar-induced subnuclear deformation. (**A**) Characterize response of cancer cells with varying malignancies to anti-metastatic drug treatment using deformation anisotropicity. (**B**) Nuclear deformation patterns and their orientation of MCF-7 cells and MDA-MB-231 cells on nanopillar arrays with or without curcumin treatment. Scale bar, 5 μm. (**C**) Anisotropy measurement of nanopillar-guided nuclear features in MCF-7 cells and MDA-MB-231 cells with or without curcumin treatment (MCF-7: N= 61 pillars (DMSO); N= 34 pillars (curcumin); MDA-MB-231: N=34 pillars (DMSO); N=52 pillars (curcumin).). (**D**) Fraction of ring deformation and line deformation in MDA-MB-231 cells and MCF-7 cells on nanopillar arrays with or without curcumin treatment (MCF-7: N= 35 cells (DMSO); N= 21 cells (curcumin); MDA-MB-231: N=27 cells (DMSO); N= 30 cells (curcumin).). Error bars represent SD. (**E**) Wound healing assay of MCF-7 cells and MDA-MB-231 cells with or without curcumin treatment for 24h. Scale bars, 200 μm. (**F**) Migration rate of MCF-7 cells and MDA-MB-231 cells under different conditions was measured using wound healing assay (N= 3 batches). Statistical significance of measurement for coherency and migration rate under different conditions was evaluated by an unpaired t-test with Welch’s correction. ****P < 0.0001; *P < 0.05; ns > 0.05.

## Discussion

Nuclear deformation on nanopillars have been reported earlier in terms of the nuclear stiffness-correlated deformation depth (*23*) and the local indentation-induced redistribution of lamin proteins (*24*). However, the reorganization of nuclear morphology, especially the pathologically related subnuclear irregular features, have been overlooked. Our study here demonstrates that, using optimized nanopillar designs, we can effectively guide the subnuclear irregularities into quantifiable patterns reflecting the cell migration ability needed for cancer metastasis. It enables quantitative evaluation of the heterogeneity in a given cell mixture and the cellular response to anti-metastatic drugs, both with single-cell resolution. This establishes an effective strategy to apply nanopillar-based technologies for cancer detection.

It is well known that the nuclear lamina formed by lamin proteins is mechanically sensitive. The extracellular forces can induce differential distribution of lamin A and B as observed in both micropipette aspiration (*36*, *37*) and nanoneedle perturbations (*24*). While intracellularly, cytoskeleton-induced nuclear membrane tension has been shown to alter the conformation of lamin A/C (*38*). In the case of nanopillars used here, it not only delivers external force to the nuclear lamina, but also triggers membrane curvature-mediated actin polymerization locally as reported earlier (*18*). Therefore, the nanopillar platform offers a unique platform to tune the subnuclear pattern of lamin A using different designs of nanopillar geometry. In addition, the formation of isotropic rings or anisotropic lines on nanopillar-deformed cancer nuclei also suggests a coordination of lamin network between adjacent nanopillars. Whether or how it correlated with tumorigenesis and cancer metastasis deserves further studies.

Alterations in both cell migration and nuclear mechanics have been found to closely correlate with cancer progression, (*39*, *40*) but how they are correlated with each other is unclear. Recent studies showed that unfolding of nuclear grooves and invaginations correlates with the enhanced cellular contractility via nuclear membrane tension and in turn strongly accelerates cell migration under whole cell compression (*41*, *42*). Similarly, we also observed that aligned subnuclear deformation correlated to increased cell migration speed. However, different from the nuclear unfolding with global compression on the entire nucleus, the alignment of subnuclear features on nanopillars are persisting but dynamically adopting different nanopillars along with cell movement. It suggests a different regulation mechanism underlying the local nuclear mechanics in response to cell migration.

The nanopillar assay described in this study enables a quantitative characterization of subnuclear deformation in cancer cells. However, the mechanism of generating subnuclear feature during cancer development is still unknown, which inevitably hinders our interpretation of the nanopillar-induced subnuclear patterns. An in-depth molecular understanding of how such irregularities evolve and whether they associate with specific genetic or metabolic alterations will greatly enrich the nanopillar-based readouts. Moreover, due to the huge morphological variations of cell nuclei among different cancer types (*1*), the nanopillar dimension optimized in this work may need to be re-evaluated with other cancer types.

## Materials and Methods

### Fabrication and characterization of nanopillar arrays

Nanopillar arrays were fabricated on the quartz chip using electron-beam lithography (EBL) and reactive ion etching (RIE). The quartz chip was cleaned with acetone and isopropyl alcohol and then spin-coated with 300 nm polymethylmethacrylate (PMMA) (MicroChem), followed by coating of one thin conductive layer, AR-PC 5090.02 (Allresist). Designed nanoscale patterns were written on the PMMA layer by electron-beam lithography (FEI Helios NanoLab 650) and the PMMA on the exposed areas was subsequently removed in the 3:1 isopropanol:methylisobutylketone solution. A Cr mask with 80 nm thickness was formed via thermal evaporation (UNIVEX 250 Benchtop), followed by lift-off with acetone. Nanopillars were finally revealed after reactive ion etching with a mixture of CF4 and CHF3 (Oxford Plasmalab 80). Characterization of nanopillar dimension was performed using SEM (FEI Helios NanoLab 650) after 10 nm chromium coating.

### Cell culture and drug treatment

Prior to cell culture, the nanostructured chips were cleaned by air plasma for 10 min and exposed to UV for 15 min. Subsequently, the nanostructured substrates were coated with fibronectin (2ug/ml, Sigma-Aldrich) for 30 minutes at 37°C. After coating, cell culture was performed on the substrates. All the cell lines used in this work were maintained in the Dulbecco’s Modified Eagle Medium (DMEM) (Gibco) supplemented with 10% fetal bovine serum (FBS) (Life Technologies) and 1% Penicillin-Streptomycin (Life Technologies) in a standard incubator at 37°C with 5% CO2. After overnight incubation and the nuclear deformation was stabilized, the MCF-7 and MDA-MB-231 cells on nanostructures are treated with curcumin (Sigma) or DMSO (Sigma). After 24-hour incubation, the treated cells and untreated cells were fixed with 4% Paraformaldehyde (PFA) Solution in PBS (Boster biological technology AR1068) for 15 minutes for subsequent immunostaining.

### Immunofluorescence staining

Cells cultured on nanopillar arrays were immunostained for lamin A or lamin B1. Cells were washed with pre-warmed PBS two times and fixed with 4% paraformaldehyde (PFA) in phosphate-buffered saline (PBS) (Boster biological technology AR1068) for 15 minutes. The cells were washed three times with PBS and then permeabilized with 0.5% Triton X-100 (Sigma) in PBS for 15 minutes. After washing with PBS for three times, samples were blocked using 5% bovine serum albumin (BSA) (Sigma) in PBS for 1 hour before staining with 1:400 anti-lamin A (Abcam ab26300) and anti-lamin B1 (gift from the Saggio lab in Sapienza University of Rome). Samples were washed three times with PBS and stained with the secondary antibody, Chicken anti-Rabbit IgG (H+L) Cross-Adsorbed Secondary Antibody, Alexa Fluor 488 (Invitrogen A21441), 1:500 in staining buffer for 1 hour under room temperature.

### Confocal imaging and live cell tracking

Imaging of the fluorescently labeled cells on nanopillar arrays was performed using laser scanning confocal microscopy (Zeiss LSM 800 with Airyscan). In particular, a Plan-Apochromat 100×/1.4 oil objective was used. During imaging, fixed cells were maintained in PBS. Z stack images were acquired with 500 nm distance between each frame. Live cell imaging and the subsequent fluorescence imaging was performed using a spinning disc confocal microscope (SDC) that is built around a Nikon Ti2 inverted microscope equipped with a Yokogawa CSU-W1 confocal spinning head, a Plan-Apo objective (100×1.45-NA), a back-illuminated sCMOS camera (Orca-Fusion; Hamamatsu). Excitation light was provided by 488-nm/150mW (Vortran) (for GFP), and all image acquisition and processing were controlled by MetaMorph (Molecular Device) software. The migration of individual cells was manually tracked using imageJ, and their migratory behavior was characterized using the method developed by a previous work (*43*).

### Transfection

For plasmid transfection in cancer cells, 1 μg plasmid was mixed with 1.5 μl Lipofectamine 3000 (Life Technologies) and 2 μl P3000 reagent (Life Technologies) in Opti-MEM (Gibco) and incubated for 20 mins at room temperature. Before the addition of the transfection mixture, cancer cells were starved with the Opti-MEM (Gibco) medium for 30 mins at 37°C. After 4 hours of incubation, the Opti-MEM (Gibco) medium was replaced with regular culture medium and the cells were allowed to recover overnight before cell sorting or live cell imaging.

### Fluorescence□activated cell sorting

Cells were sorted by using the BD FACS Aria II, and gating was done using the BD FACSDiva™ software (Becton, Dickinson Biosciences). Dead cells were excluded from analysis on the basis of FSC/SSC; cell aggregates or small debris were excluded from analysis on the basis of side scatter (measuring cell granularity) and forward scatter (measuring cell size); lastly, GFP positive cells were sorted on the basis of fluorescence intensity.

### Wound healing assay

Cells were maintained in 35 mm dishes for each cell line until approximately 90% confluent. Scratch was made in the confluent monolayer of cells with a sterile 200-μl pipette tip, and fresh culture medium was replaced. Brightfield microscopic pictures were taken of the same field at 24 hours. Migration rate was measured by quantifying the closure area within the same time frame using ImageJ.

### Statistical analysis

Welch’s t tests (unpaired, 2 tailed, not assuming equal SD) were used to evaluate the significance. All tests were performed using Prism (GraphPad Software). Data are presented as mean ± SEM or mean ± SD as stated in the figure captions. All experiments were repeated at least twice, unless otherwise explicitly stated in the figure captions.

## Author contribution

Y.P.Z. and W.Z. conceived the idea and designed the experiments. Y.P.Z. and Y.Y.Z. performed most of the experiments and data analysis. Y.P.Z. and W.G. performed fabrication and SEM experiments. P.C., I.S. and G.V.S. provided material and experimental supports. Y.P.Z. and W.Z. drafted the manuscript. W.G., P.C., I.S. and G.V.S. edited the manuscript. All of the authors discussed and commented on the manuscript.

## Data and materials availability

All data needed to evaluate the conclusions in the paper are present in the paper and/or the Supplementary Materials.

