## Supplementary Info for "Guiding Irregular Nuclear Morphology on Nanopillar Array for Malignancy Differentiation in Tumor cells"

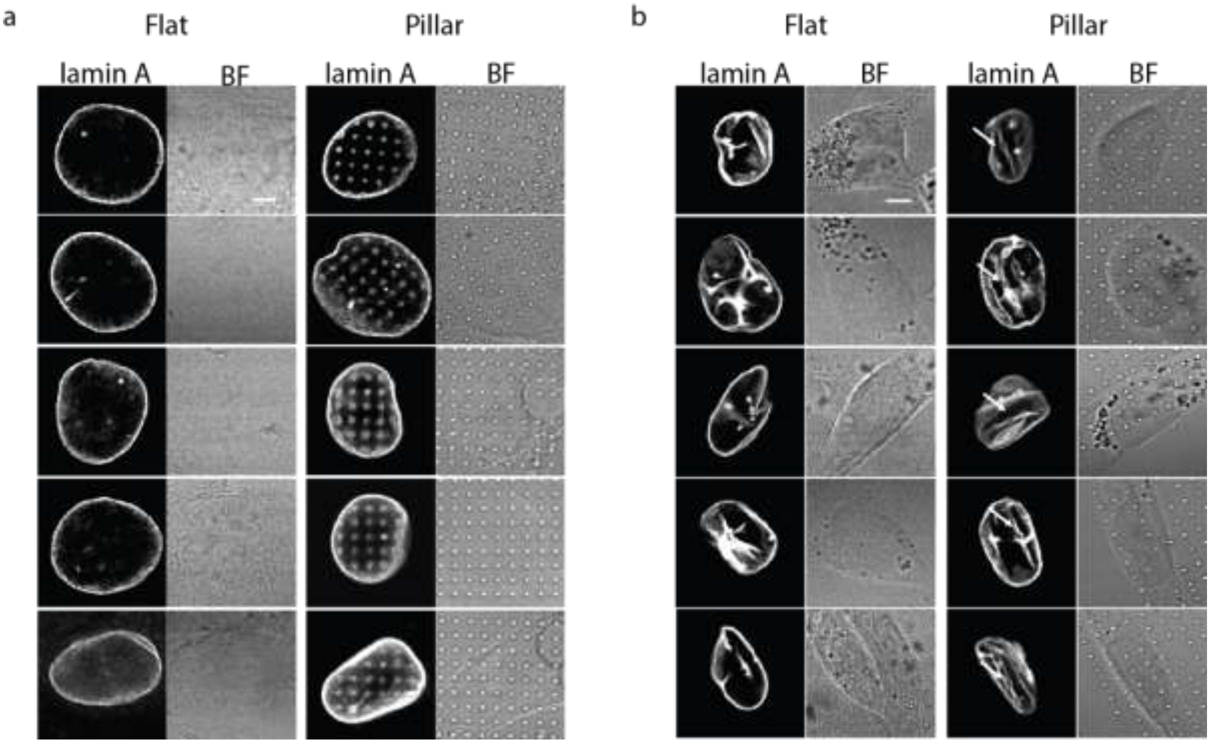


**Figure S1. Representative images of nuclear morphology of MDA-MB-231 cells and MCF-7 cells on flat versus on nanopillar arrays. a)** Nuclear morphology of MCF-7 cells on flat surfaces and nanopillar-guided subnuclear features in MCF-7 cells. **b)** Nuclear morphology of MDA-MB-231 cells on flat surfaces and nanopillar-induced subnuclear shape irregularities in MDA-MB-231 cells. Scale bars, 5 µm.

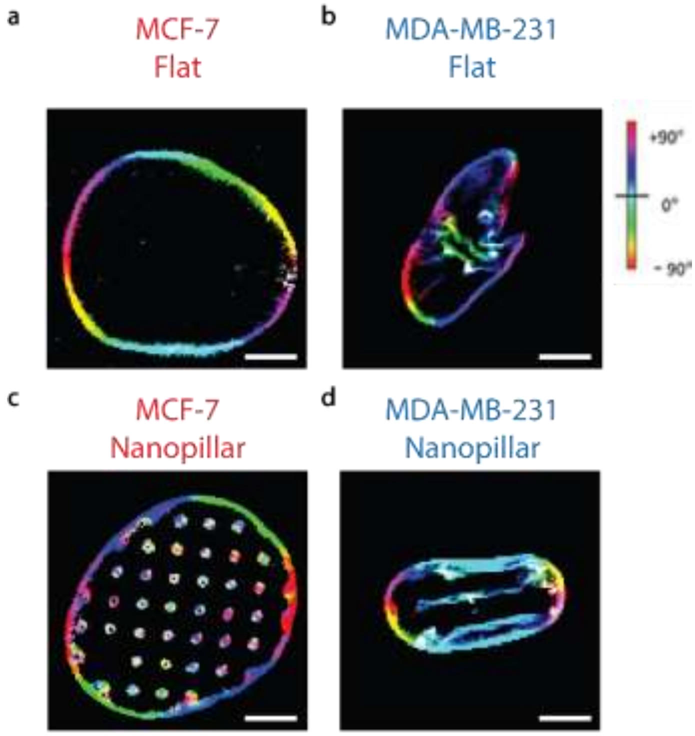


**Figure S2. Color representation of oriented nuclear morphology of MCF-7 cells and MDA-MB-231 cells on flat surfaces and nanopillar arrays.** For MCF-7 cells, all-angle-oriented features with multiple color evenly displayed in MCF-7 cells on both flat **(a)** and nanopillar arrays **(c)**. For MDA-MB-231 cells, random orientation of nuclear shape irregularities showed on flat surfaces in evenly displayed colors (**b)**, while highly aligned features on nanopillar arrays exhibit dominant light blue **(d)**. Scale bars, 5 μm.

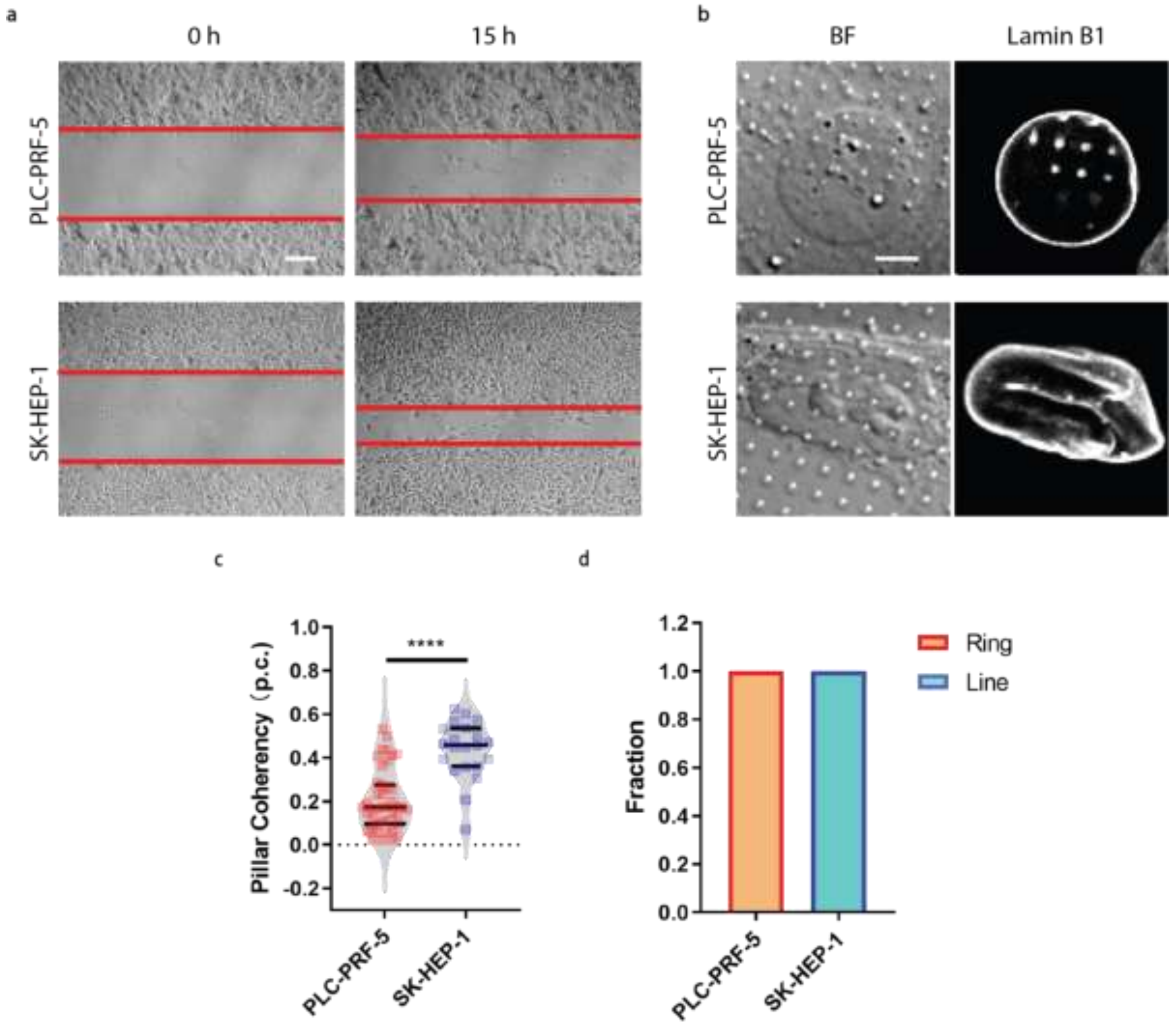


**Figure S3. Correlation of naopillar-guided anisotropicity and malignancy in liver cancer cells. a)** Wound healing assay revealed liver cancer cell lines with varying motility. Scale bar, 100 µm. **b)** Nanopillar-guided nuclear features in PLC-PRF-5 and SK-HEP-1 cells. Scale bar, 5 µm. **c)** Pillar coherency measurement of nanopillar-guided nuclear features in different liver cancer cells. **d)** Fraction of cells with classified ring or line deformation in different liver cancer cells. Statistical significance of measurement for coherency under different conditions was evaluated by an unpaired t-test with Welch's correction. ****$P < 0.0001$.

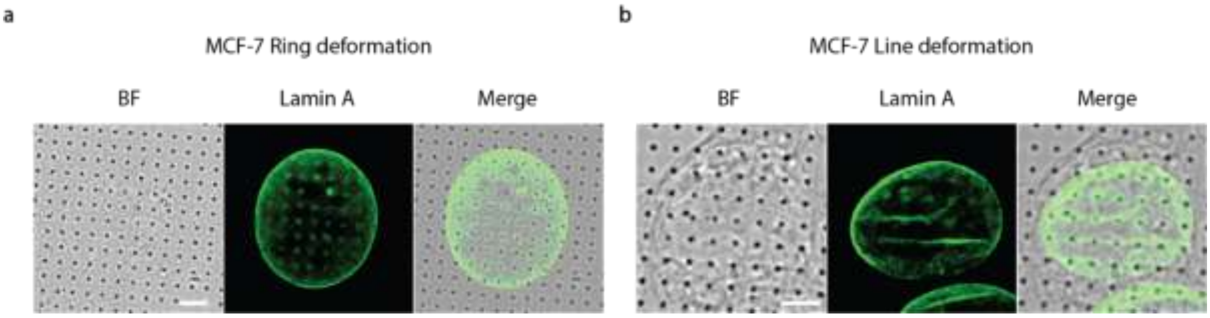


**Figure S4. MCF-7 cells with (a) ring and (b) line deformation of nucleus on nanopillars**. Scale bars, 5 μm.

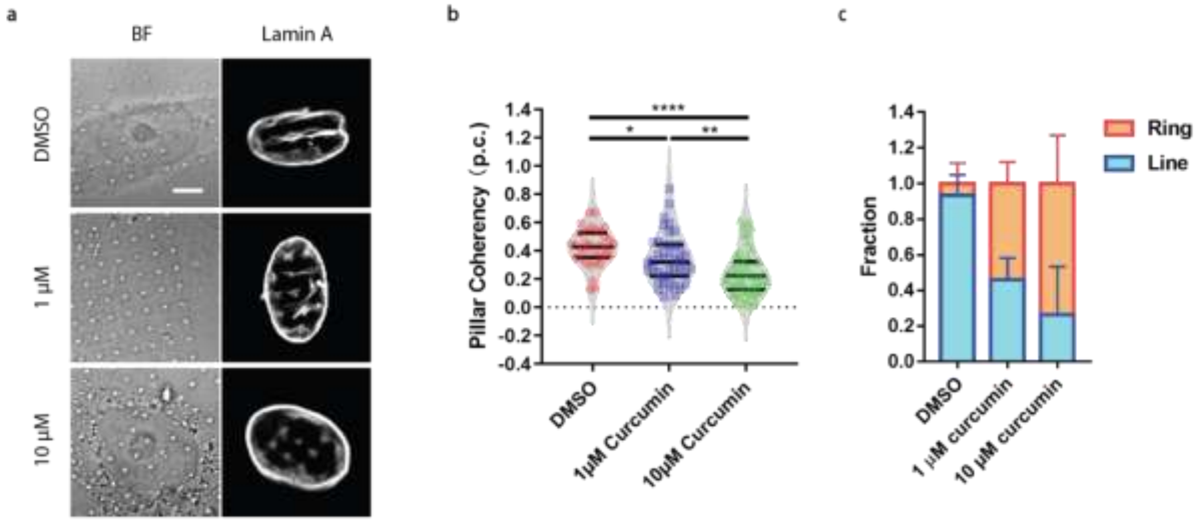


**Figure S5. Nanopilalr-guided subnuclear anisotropicity correlates with the concentration of anti-metastatic drug, curcumin. a)** Nanopillar-guided nuclear features in MDA-MB-231 cells with DMSO, 1 μM and 10 μM curcumin treatment. Scale bar, 5 μm. **b)** Pillar coherency measurement of nanopillar-guided nuclear features in cells under the curcumin treatment with different concentrations. **c)** Fraction of cells with classified ring or line deformation in MDA-MB-231 cells with DMSO, 1 μM and 10 μM curcumin treatment. Statistical significance of measurement for coherency under different conditions was evaluated by an unpaired t-test with Welch's correction. ****$P < 0.0001$;**$P < 0.01$; *$P < 0.05$.

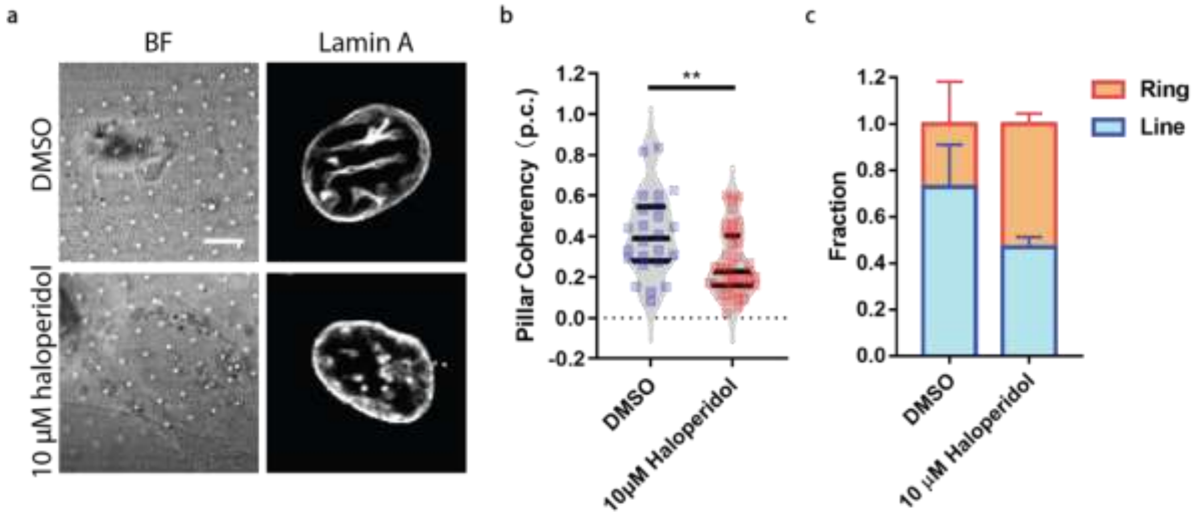


**Figure S6. Effect of anti-metastatic drug, haloperidol, on nanopillar-guided subnuclear deformation in MDA-MB-231 cells. a)** Nanopillar-guided nuclear features in MDA-MB-231 cells with or without 10 μM haloperidol treatment. Scale bar, 5 μm. **b)** Pillar coherency measurement of nanopillar-guided nuclear features measured with DMSO and haloperidol treatment. **c)** Fraction of cells classified with ring or line deformation in MDA-MB-231 cells with or without 10 μM haloperidol treatment. Statistical significance of measurement for coherency under different conditions was evaluated by an unpaired t-test with Welch's correction. $^{**}P < 0.01$.
